## Supplementary Figures and Tables for "Quantitative 3D Imaging of the Cranial Microvascular Environment at Single-Cell Resolution"

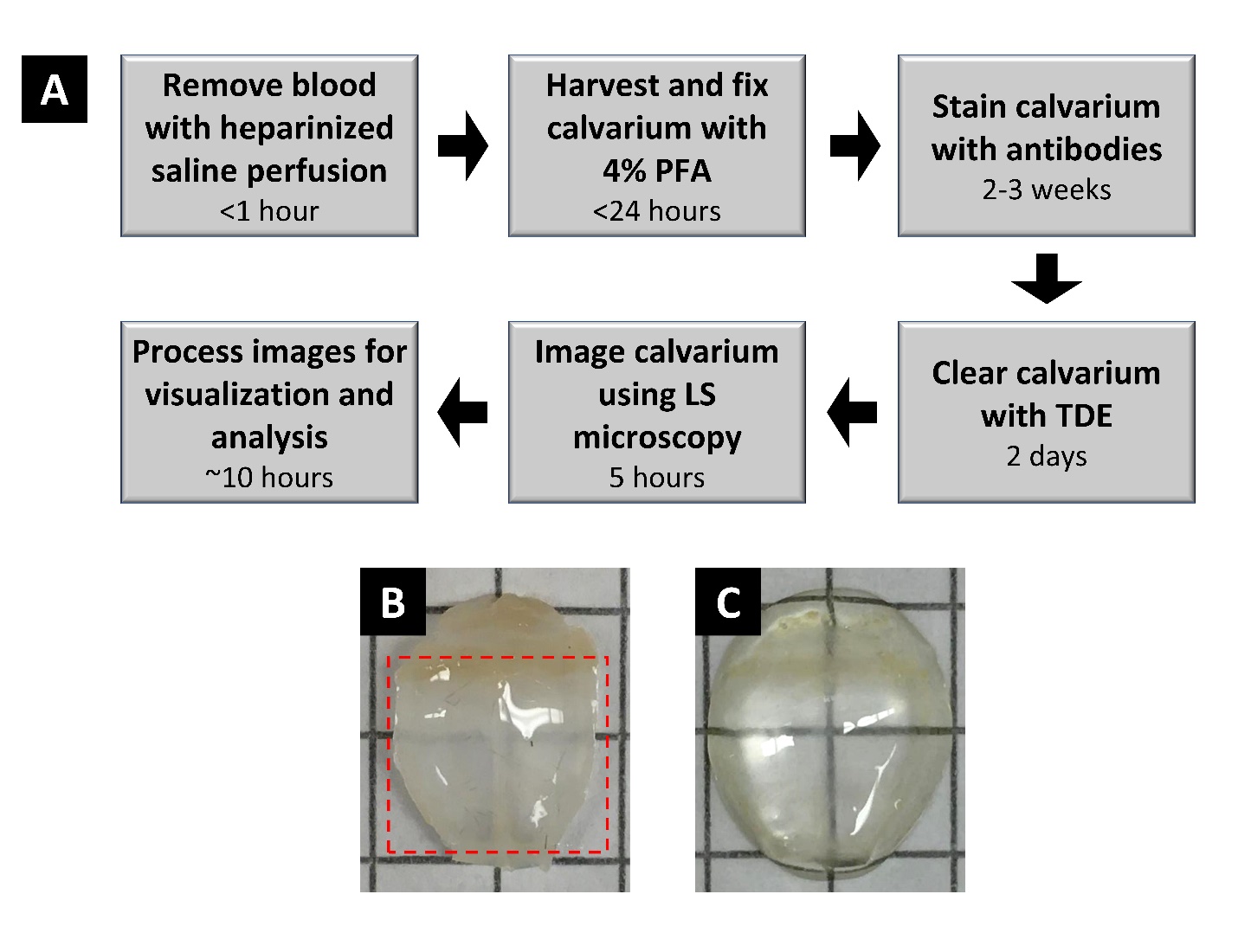

**Supplemental Figure 1. Quantitative 3D light-sheet imaging pipeline.** A) Diagram displaying the steps to harvest, stain, clear, and image calvaria for data analysis. The entire procedure—including quantitative analysis—takes 3-4 weeks to complete. B) Non-decalcified calvarium before optical clearing. The dotted red lines indicate the region of the calvarium that was imaged. C) Non-calcified calvarium following clearing with TDE.

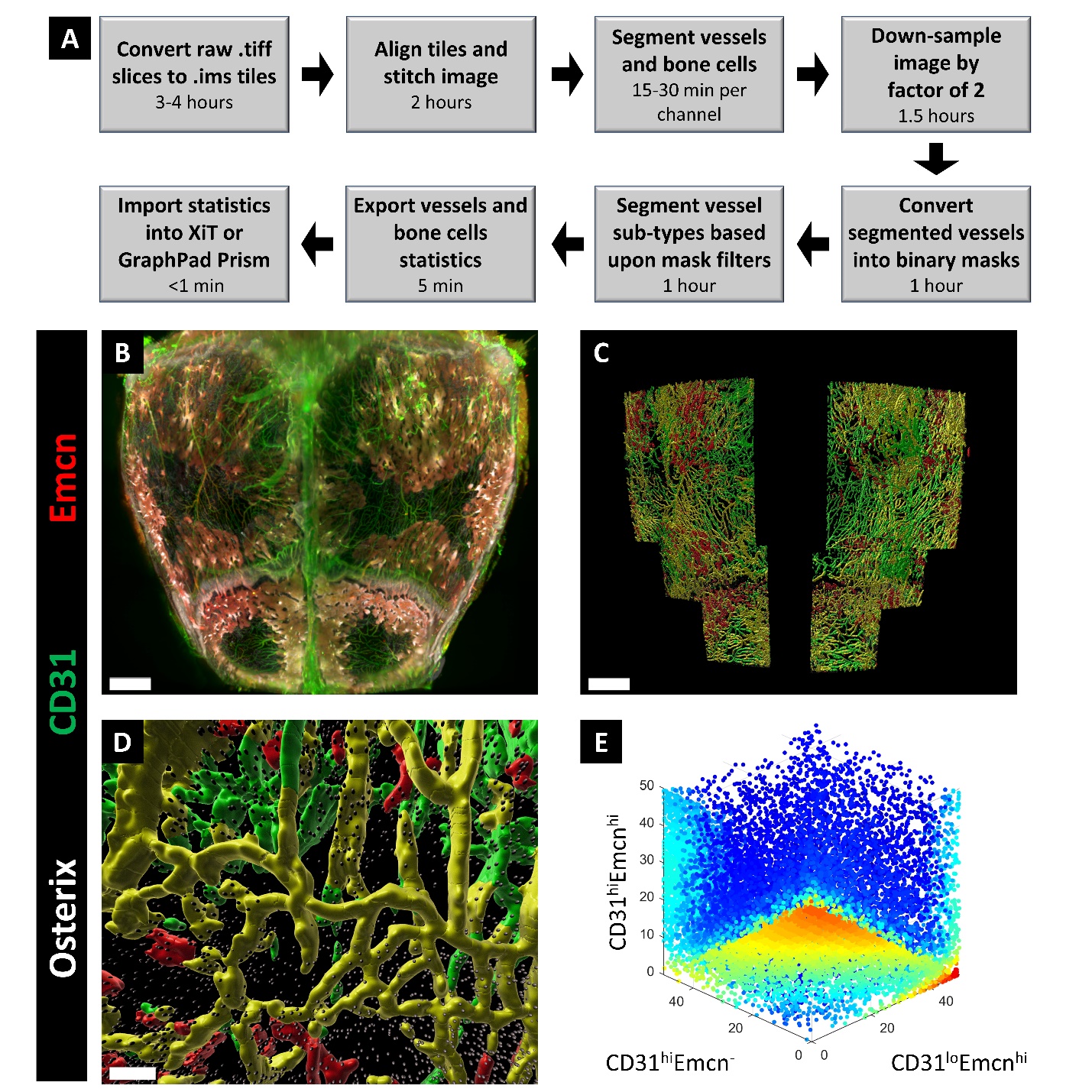

**Supplemental Figure 2. Quantitative analysis pipeline for characterizing calvarial vessel phenotype and skeletal progenitor distribution in 3D.** A) Diagram displaying the steps performed to analyze each dataset. The entire pipeline takes approximately 10 hours to complete using a high-performance workstation. B) 3D projection of calvarium stained for CD31, Emcn, and Osterix. C) 3D segmentation of calvarial vessel phenotypes and Osterix+ cells in volume of interest (sagittal suture not included due to interference of sagittal sinus with the segmentation). D) Zoomed-in portion of (C) displaying the spatial relationships of Osterix+ cells to each vessel phenotype. Segmented colors represent the following: CD31^hi^Emcn^-^ vessels (green), CD31^hi^Emcn^hi^ vessels (yellow), CD31^lo^Emcn^hi^ vessels (red), Osterix+ cells (gray). E) 3D plot representing the distance of individual Osterix+ cells to each vessel phenotype from (C). Color represents density of cells relative to regions of the plot. Scale bars: 1000 μm (B,C); 200 μm (D)

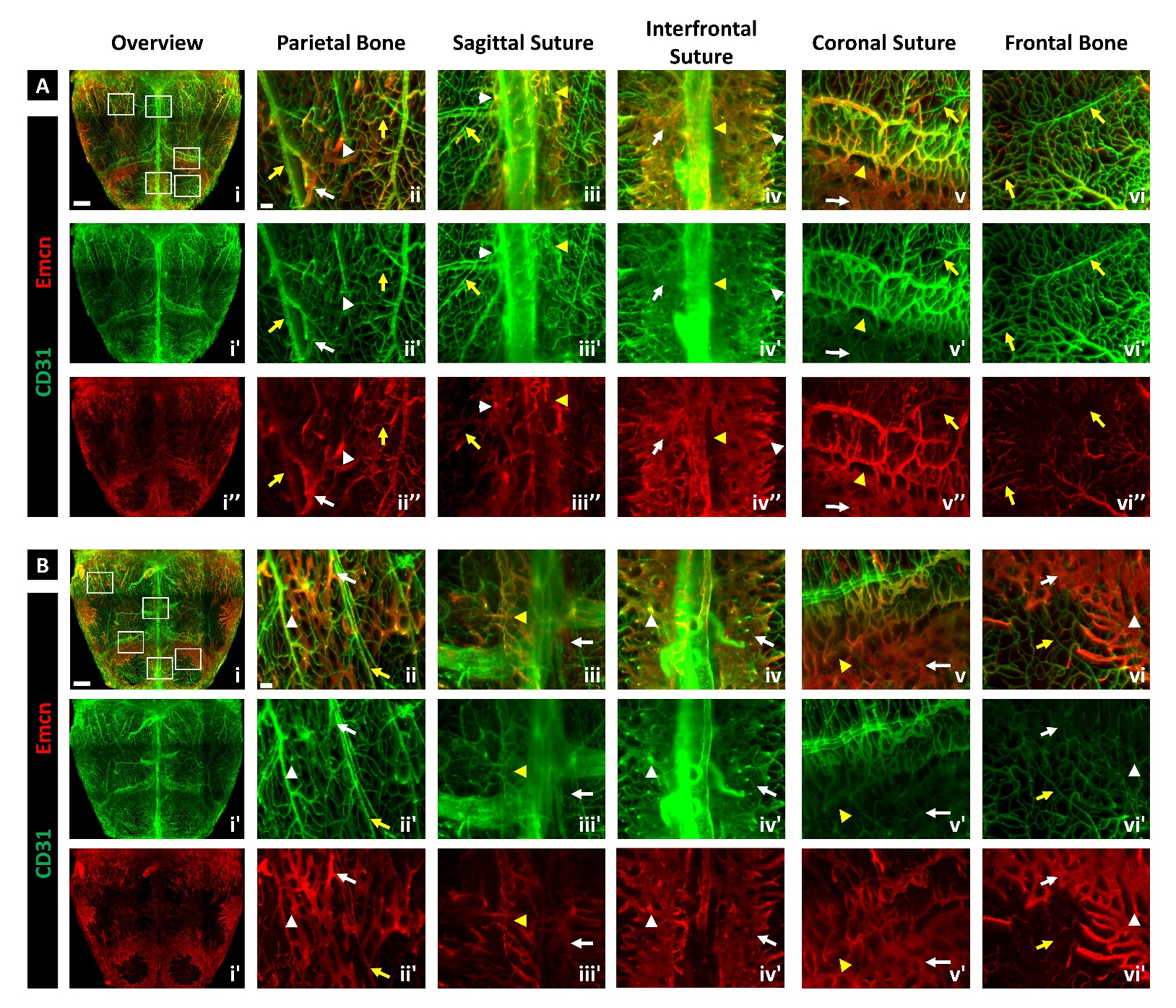

**Supplemental Figure 3. 3D map of vessel phenotypes in calvaria of 4-week-old mice.** A-B) MIP of vessel phenotypes in two independent calvaria, as originally depicted in Figure 2. Aii-vi and Bii-vi represent different regions of the calvarium marked in Ai and Bi. CD31^hi^Emcn^-^ and CD31^hi^Emcn^hi^ vessels in the periosteum and dura mater (yellow arrows) connect to CD31^hi^Emcn^hi^ and CD31^lo^Emcn^hi^ marrow vessels via CD31^hi^Emcn^hi^ transcortical vessels (white arrowheads). Vessels in the periosteum are prevalent across the parietal and frontal bones, while marrow vessels are restricted to regions nearby the sutures (yellow arrowheads). Scale bars: 1000 μm (Ai, Bi), 100 μm (Aii-v, Bii-v)

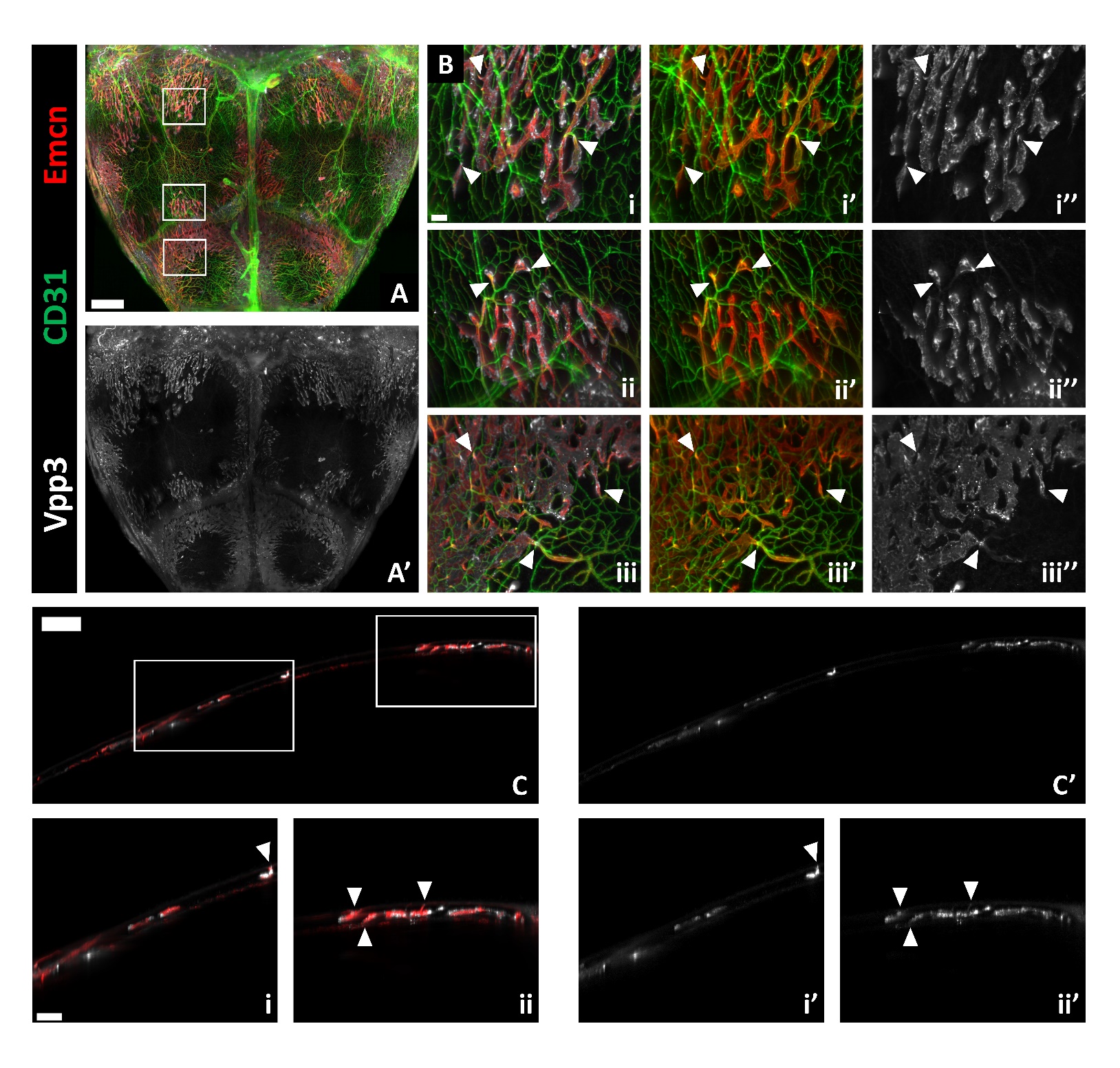

**Supplemental Figure 4. Osteoclasts reside in marrow cavities and transcortical canals.** A) MIP of vessels and Vpp3+ osteoclasts in the calvarium from a 4-week-old mouse. B) Zoomed-in regions from (A) demonstrating spatial distribution of osteoclasts relative to vessels. Arrowheads point to regions where osteoclasts reside at or adjacent to transcortical canals. C) 40 μm-thick sagittal section displaying the distribution of osteoclasts relative to Emcn^hi^ transcortical and marrow vessels. Arrowheads indicate regions where osteoclasts reside nearby Emcn^hi^ transcortical vessels. Scale bars: 1000 μm (A); 500 μm (C); 200 μm (Ci-ii); 100 μm (B)

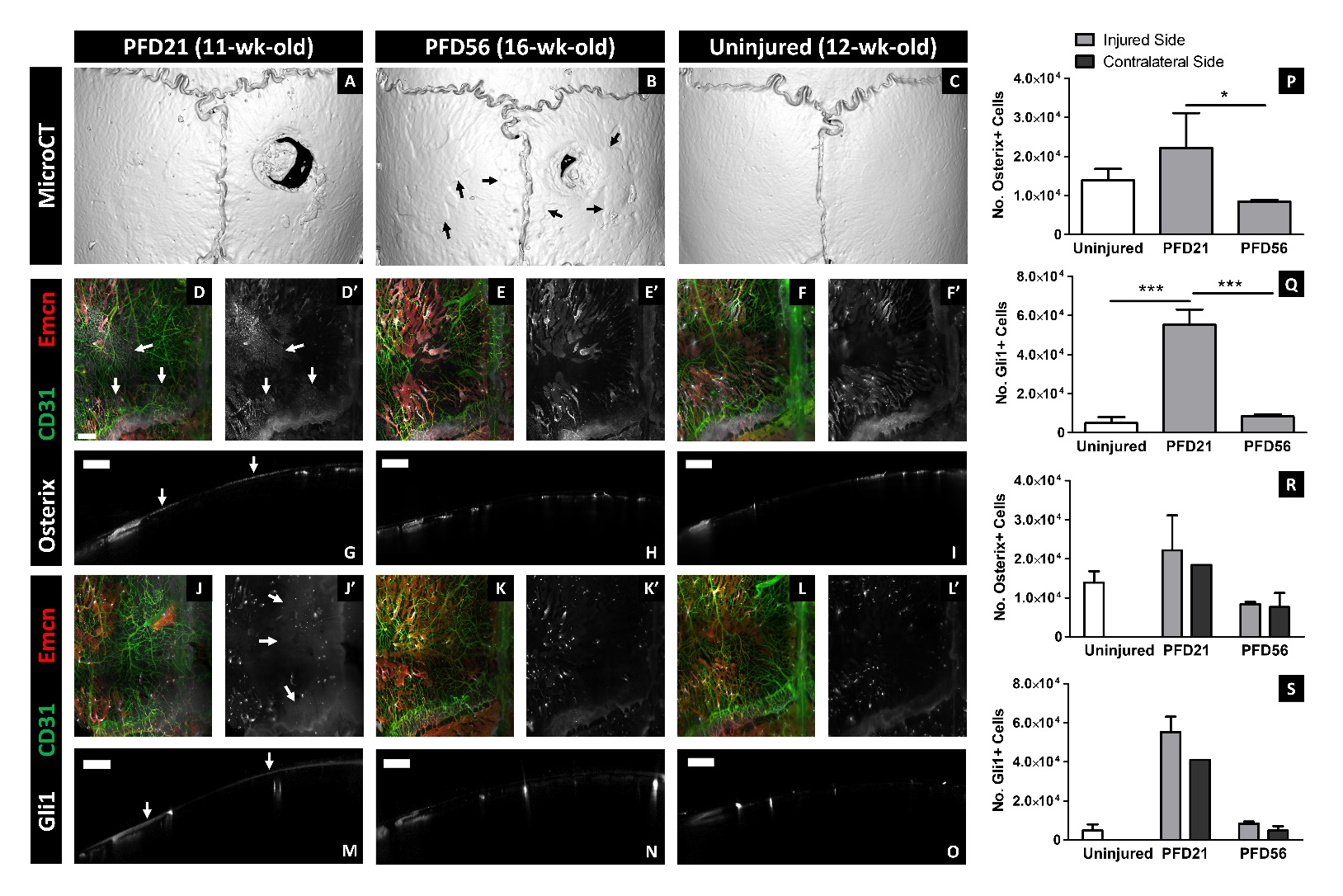

**Supplemental Figure 5. Localized calvarial injury stimulates systemic skeletal progenitor expansion.** A-C) MicroCT 3D surface rendering of calvaria at PFD21 (A) and PFD56 (B) compared to an uninjured calvarium (C). Black arrows point to regions of excess mineral deposition along the cortical surface of injured calvaria (B). D-F) MIP of calvarial vessels and Osterix+ skeletal progenitors in the contralateral parietal bone at PFD21 (D) and PFD56 (E) compared to an uninjured control (F). G-I) 40 μm-thick sagittal sections from (D-F) displaying distribution of Osterix+ progenitors along the thickness of the contralateral parietal bone. J-L) MIP of calvarial vessels and Gli1+ skeletal progenitors in the contralateral parietal bone at PFD21 (J) and PFD56 (K) compared to an uninjured control (L). M-O) 40 μm-thick sagittal sections from (J-K) displaying distribution of Gli1+ progenitors along the thickness of the contralateral parietal bone. White arrows mark regions of skeletal progenitor expansion within the periosteum at PFD21 (D, G, J, M). P-Q) Quantification of skeletal progenitors in the ipsilateral side of the injured parietal bone compared to the uninjured control (n=3). R-S) Quantification of skeletal progenitors in the ipsilateral and contralateral side of the injured parietal bone compared to the uninjured control (n=3 for control, ipsilateral PFD21, ipsilateral and contralateral PFD56; n=1 for contralateral PFD21). Data are mean ± SD. Statistics were performed using a one-way ANOVA with Tukey post-hoc test (P, Q). ***p<0.001, **p<0.01, *p<0.05 where designated. Scale bars: 500 μm (D-O)

**Supplemental Video 1. 3D projection of blood vessels and Osterix+ skeletal progenitors in the murine calvarium.** Fluorescent labels are denoted using the following pseudo colors: green (CD31), red (Emcn), and gray (Osterix). Segmentation is shown for the following: CD31^hi^Emcn^-^ arteries and arterioles (green), CD31^hi^Emcn^hi^ capillaries (gold), CD31^lo^Emcn^hi^ sinusoids (red), and Osterix+ skeletal progenitors (gray).

Video included as a separate file.

**Supplemental Table 1.** List of all key reagents and resources used in this study.

| **Reagent or resource** | **Source** | **Identifier** |
| --- | --- | --- |
| ***Antibodies*** | | |
| Goat anti-mouse/rat CD31 (1:100) | R&D Systems | AF3628 |
| Rat anti-mouse/rat Endomucin (1:50) | Santa Cruz Biotechnology | sc-65495 |
| Rabbit anti-mouse/rat/human Osterix (1:200) | Abcam | ab209484 |
| Rabbit anti-mouse/human Gli1 (1:100) | Sigma Aldrich | SAB4301901-100UL |
| Rabbit anti-mouse/rat/human Vpp3 (ATP6V1B1 + ATP6V1B2; 1:200) | Abcam | ab200839 |
| Donkey anti-goat AF800 plus, 0.67 mg/mL, (1:50) | Thermo Fisher Scientific | A32930 |
| Donkey anti-rabbit AF647 plus, 0.67 mg/mL (1:150) | Thermo Fisher Scientific | A32795 |
| Donkey anti-rat biotin, 0.75 mg/mL (1:100) | Thermo Fisher Scientific | A18749 |
| Streptavidin AF555 conjugate, 0.67 mg/mL (1:100) | Thermo Fisher Scientific | S32355 |
| ***Reagents*** | | |
| Heparin sodium salt from porcine mucosa | Sigma Aldrich | H3393-50KU |
| Paraformaldehyde, 16% aq. soln., methanol free | Alfa Aesar | 433689M |
| Normal donkey serum | Sigma Aldrich | D9663-10ML |
| Trizma base | Sigma Aldrich | T6066-1KG |
| Trizma hydrochloride | Sigma Aldrich | T5941-1KG |
| Sodium chloride | Sigma Aldrich | S5886 |
| Tween 20 | Sigma Aldrich | P7949 |
| Dimethylsulfoxide | Thermo Fisher Scientific | PI20688 |
| Streptavidin/Biotin Blocking Kit | Vector Laboratories | SP-2002 |
| 2,2-thiodiethanol | Sigma Aldrich | 166782-500G |
| ***Animal drugs and materials*** | | |
| Ketamine HCl (100 mg/mL) | VetOne | 501072 |
| Xylazine Injection (20 mg/mL) | Akorn, Inc. | N/A |
| pTH (1-34) | Bachem | 4011474 |
| Povidone-Iodine Swabsticks | Fisher Scientific | 06-669-83 |
| 6-0 nylon monofilament sutures | Ethicon | 1665G |
| Buprenorphine SR-LAB (0.5 mg/mL) | ZooPharm | N/A |
| Ideal Micro-Drill | Harvard Apparatus | 72-6065 |
| 1 mm carbide inverted cone burr | Roboz Surgical Instrument Co. | RS-6282C-35 |
| Stereotactic frame (ask Alex for info) | KOPF, David Kopf Instruments | 900LS |
| Surgical microscope (ask Alex for info) | Carl Zeiss | NC-4 |
| ***Instruments and hardware*** | | |
| Ultramicroscope II, 2x zoom body configuration | Miltenyi Biotec (formerly manufactured by LaVision Biotec) | N/A |
| LVMI-Fluor 2x objective lens with dipping cap (0.5 NA, 1.33-1.57 RI range, 5.6 mm WD) | Miltenyi Biotec (formerly manufactured by LaVision Biotec) | N/A |
| Dell Precision 7820 Tower | Dell | T7820X |
| Portable SSD T5 USB 3.1 2 TB | Samsung | MU-PA2T0B |
| Skyscan 1275 μCT | Bruker | N/A |
| ***Software*** | | |
| Imaris 9.5 Single Full with ClearView plus Stitcher | Bitplane Inc. | N/A |
| XiT | See ref. [14] | N/A |
| CTAN and CTVOL | Bruker | N/A |
| Mimics 14 | Materialise |  |
| Microsoft Excel 2019 | Microsoft | N/A |
| GraphPad Prism 5 | GraphPad Software | N/A |
